# EHMT2 Controls Neural Crest-Derived Craniofacial Development but is Dispensable in Limb Development

**DOI:** 10.1101/2025.09.07.674526

**Authors:** Ye Liu, Yaguang Zhao, Minmin Liu, Paul Kim, Ji Liao, Di Lu, Huadie Liu, Piroska E. Szabó, Tao Yang

## Abstract

Post-translational modifications of histones, such as methylation of histone H3 at lysine 9 (H3K9), play critical roles in regulating chromatin structure and gene expression. EHMT2 (also called G9A), a histone methyltransferase, mediates H3K9 mono- and dimethylation and has been implicated in both transcriptional repression and context-specific gene activation. Although global knockout of the mouse *Ehmt2* gene results in early embryonic lethality, tissue-specific knockouts have uncovered diverse roles in organ development. However, how EHMT2 contributes to skeletal development in a lineage-specific manner remains to be fully elucidated. Here, we investigated the role of EHMT2 in skeletal development by conditionally inactivating *Ehmt2* in neural crest-derived and mesoderm-derived progenitors using *Wnt1-Cre* and *Prx1-Cre* mouse lines, respectively. Loss of *Ehmt2* function in neural crest cells led to postnatal growth failure and craniofacial defects, including delayed intramembranous ossification and malformations of the jaw and cranial base. Transcriptomic analysis of neural crest cells revealed disrupted chromatin regulatory networks, reduced expression of proliferation-associated genes, and upregulation of inflammatory pathways. In contrast, inactivation of *Ehmt2* in *Prx1*-expressing mesodermal progenitors had minimal impact on limb and cranial bone development, with no significant alterations in bone mass or osteoblast function. Together, these results reveal a lineage-specific requirement for EHMT2 in neural crest-derived skeletal tissues, suggesting that distinct progenitor populations exhibit differential dependency for bone development.

## Introduction

Chromatin accessibility and gene expression in eukaryotes are regulated by post-translational modifications of histone proteins. Among these, methylation of histone H3 at lysine 9 (H3K9) is a key repressive mark that helps establish and maintain transcriptional silencing. This modification is catalyzed by several SET domain-containing histone methyltransferases (KMTs), including SUV39H1/SUV39H2, SETDB1, and EHMT1/2 (also known as GLP/G9A) ^1^. Suv39h1/2 and Setdb1 catalyze H3K9 trimethylation (H3K9me3) at pericentric heterochromatin and transposable elements, whereas EHMT2 and EHMT1mediate H3K9 mono- and dimethylation (H3K9me1/2) in euchromatic regions ^2,3^.

In addition to its canonical repressive functions, EHMT2 has been shown to play non-canonical roles in gene regulation, including supporting transcription of certain genes in a context-dependent manner.

EHMT2 and EHMT1 usually function as a heteromeric complex and are both essential for mouse embryogenesis; individual knockouts result in early lethality and global reduction of H3K9me1/2 ^3^. Conditional knockout studies have uncovered tissue-specific roles for *Ehmt2* in organ development, including the heart, adipose tissue, and bone. For instance, *Ehmt2* deletion in adipose tissue increases pro-apoptotic gene expression and alters tissue mass ^4^, and loss of *Ehmt2* in cardiomyocytes disrupts cardiac homeostasis and induces hypertrophy in a context-dependent manner ^5^. In contrast, *Ehmt2* appears dispensable for skeletal muscle development ^6^.

*Ehmt2* has been implicated in the regulation of cranial bone formation ^7^, but its lineage-specific roles in skeletal development remain incompletely understood. In mammals, craniofacial bones are largely generated through intramembranous ossification by neural crest-derived progenitors, whereas the axial and appendicular skeleton arises from mesoderm-derived osteochondral progenitor cells primarily through endochondral ossification, and some flat bone may develop through intramembranous ossification. Neural crest cells (NCCs) are a transient, multipotent cell population that emerges from the neural plate border during early embryogenesis ^8^. They migrate extensively and differentiate into diverse cell types, including peripheral neurons, glia, melanocytes, and craniofacial skeletal components. Due to their remarkable plasticity in tissue development, NCCs have been described as the “fourth germ layer” ^9^. In contrast, osteochondral progenitors, which originated from mesoderm, contribute primarily to limb and torso skeleton development ^10^. Recent clinical studies have identified a loss-of-function mutation in human *EHMT2* as a cause of neurodevelopmental defects with craniofacial dysmorphisms, implicating its essential role in human craniofacial development ^11^. Another study described a de novo missense mutation in the catalytic SET domain of *EHMT2*, leading to reduced H3K9 methyltransferase activity and a Kleefstra syndrome-like phenotype ^12^. These case reports suggest that EHMT2 is involved in neurodevelopmental disorders with craniofacial abnormalities, although the underlying mechanisms remain largely unknown.

To examine the lineage-specific role of *Ehmt2* in skeletal development, we conditionally inactivated *Ehmt2* using *Wnt1-Cre* and *Prx1-Cre* drivers to target neural crest- and mesoderm-derived lineages, respectively ^13,14^. Loss of *Ehmt2* in neural crest cells (*Wnt1-Cre*) led to a delay in craniofacial bone formation, while the facial and limb skeletons of *Prx1-Cre* mice remained unaffected. These findings highlight a lineage-dependent role for *Ehmt2* in the development of the skeletal system based on the origins of corresponding progenitors.

### Materials and Methods Mice and tissue collection

All animal experiments were performed according to American Association for Accreditation of Laboratory Animal Care (AALAC), with Institutional Care and Use Committee-approved protocols at Van Andel Institute. The conditional mutant *Ehmt2* flox/flox (*Ehmt2*^*f/f*^) mice line, which carried a floxed SET domain, was previously generated by Dr. Piroska E. Szabó’s lab on a 129S1/ SvImJ genetic background^15^. *Wnt1-Cre* mice and *Prx1-Cre* mice were obtained from Jackson Lab. To generate tissue-specific knockouts, *Ehmt2*^*f/f*^ mice were crossed with *Wnt1-Cre* mice or *Prx1-Cre* mice, resulting in deletion of the floxed SET domain in the Cre-expressing lineages. The mice were maintained in a C57BL/6 background, fed with a normal diet (LabDiet 5021, Purina, St. Louis, MO, USA). The left femurs were used for µCT scanning, and the right femurs were fixed in 10% formalin for 48 hours and embedded in methyl methacrylate resin (MMA) or in paraffin after demineralization (10% EDTA, 14 days) for bone histomorphometry analysis. The µCT and bone histomorphometry data were quantified in a blinded manner.

### Microcomputed Tomography (µCT)

Mouse femurs and skulls were collected and scanned using the SkyScan1172 µCT imaging system (Bruker µCT, Kontich, Belgium) at a voxel size of 13.3 µm (59 kV, 167 µA). Imaging was performed according to the guidelines recommended by the Journal of Bone and Mineral Research (JBMR) ^16^.

Cross-sectional slices were reconstructed with NRecon software (SkyScan), and bone parameters were calculated using CTAn software (SkyScan, Bruker µCT, Belgium). 3D images were visualized and analyzed with CTVol software (SkyScan, Bruker µCT, Belgium). The following bone parameters were calculated: bone volume/total volume (BV/TV), trabecular number (Tb.N), trabecular thickness (Tb.Th), trabecular separation (Tb.Sp), cortical bone thickness (Cs.Th), cortical bone area (B.Ar), total cross-sectional area (T.Ar), and the ratio of cortical area to total area (B.Ar/T.Ar).

### RNA-seq and data processing

NCCs were isolated from the first branchial arch of E10.5 embryos from (*Ehmt2*^*f/f*^) WT and *Wnt1-Cre*; *Ehmt2*^*f/f*^ (KO) mice (n=2 per group), as previously described ^17^. Total RNA was isolated by Qiagen RNeasy Plus Micro kit (Cat. No. 74034). RNA sequencing was performed by the Genomics Core at Van Andel Institute. Approximately 40 million reads were generated per library. Paired-end reads were quality-trimmed using *Trim Galore* (v0.6.7) with default parameters to remove adapter sequences and low-quality bases (Phred score < 20). *FastQC* (v0.11.9) was run in parallel to assess read quality before and after trimming. Trimmed reads were aligned to the mouse reference genome (mm39) using *HISAT2* (v2.2.1) with the --rna-strandness RF parameter to account for strand-specificity. Alignments were converted to BAM format, sorted, and indexed using *samtools* (v1.12). Duplicate reads were not removed to retain transcript diversity information. Read counts per gene were generated using *featureCounts* (v2.0.3) from the Subread package. Aligned reads were assigned to exons in the GENCODE vM30 annotation (mm39) with the following parameters: -t exon -g gene_name. Count matrices were extracted for downstream analysis, in R (V4.4.2) with *DESeq2* (v1.40.2) for differential gene expression. Genes with |log2FC| > 1 and *p* < 0.05 were considered significant. Euclidean distances between samples were calculated from variance-stabilized transformed (VST) counts (batch-corrected via limma::removeBatchEffect) and visualized by hierarchical clustering (pheatmap). PCA is conducted on the top 500 most variable genes using *DESeq2::plotPCA* (modified to incorporate batch correction). Gene Set Enrichment Analysis (GSEA) was performed using the fgsea package (v1.24.0) with gene sets from the MSigDB Hallmark collection (c2.all.v2024.1) and Reactome pathways (c2.cp.reactome.v2024.1). Pathways with an unadjusted p-value < 0.05 were considered significant. The top enriched pathways were ranked by normalized enrichment score (NES). Volcano plot was generated with *EnhancedVolcano* (|log2FC| > 1, *p* < 0.05). Normalized counts for NCC regulators (e.g., *Meis1, Zeb1*) were plotted as barplots (mean ± SD) with custom sex-matched colors.

### Bone histomorphometry

Femurs from 2-month-old *Prx1-Cre*; *Ehmt2*^*f/f*^ and *Ehmt2*^*f/f*^ (control) mice (n = 15; 8 males and 7 females) were collected for bone formation analyses. Mice were injected intraperitoneally with calcein green (10 mg/kg, Sigma, Cat# C0875) at 12 days, followed by alizarin complexone (20 mg/kg, Sigma, Cat# A3882) at 5 days before sacrifice. After euthanasia, femurs were fixed in 4% formalin for 48 hours and then embedded in methyl methacrylate. Sections were cut at 7–10 µm thickness, mounted on slides, and imaged using a Zeiss Axio 2 microscope (Thornwood, NY, USA). Bone formation parameters, including mineral apposition rate (MAR), bone formation rate/bone surface (BFR/BS), and mineral surface/bone surface (MS/BS), were calculated using BioQuant Osteo software (BioQuant, Nashville, TN, USA).

### Mouse bone marrow stromal cell culture and osteoblast differentiation assay

Bone marrow was flushed from the tibiae and femurs of mice using α-MEM supplemented with 10% fetal bovine serum (FBS), 1% penicillin-streptomycin (P/S), and 1% L-glutamine. The collected bone marrow cells were cultured in the same medium. After 48 hours, non-adherent cells were removed by washing with PBS, and the adherent bone marrow stromal cells (BMSCs) were maintained with medium changes every two days. Once cells reached confluency, they were trypsinized and seeded in triplicate at 7.5 × 10^4^ cells per well in 24-well plates containing complete α-MEM. After 1–2 days (defined as Day 0), the medium was replaced with osteogenic differentiation medium composed of α-MEM containing 10% FBS, 1% P/S, 100 nM dexamethasone (from a 1 mM stock, 10,000× dilution), 10 mM β-glycerophosphate (from a 1 M stock, 100× dilution), and 0.05 mM L-ascorbic acid-2-phosphate (from a 10 mM stock, 200× dilution).

The differentiation medium was replaced every two days. Alkaline phosphatase (ALP) staining was performed on Day 4, and Alizarin Red staining was conducted on Days 8 and 14.

### Total RNA isolation and qRT-PCR

Total RNA from the tibia and femur bones was extracted using TRIzol reagent (Invitrogen) by following the standard protocol. 500 ng of RNA were subjected to the synthesis of first-strand cDNA using a SuperScript^TM^ VILO^TM^ cDNA Synthesis Kit (Invitrogen, 11754050). Quantitative PCR (qPCR) was performed on a StepOne PCR instrument using SRYBR Green QPCR Master Mix (Invitrogen, 4472908). Primers used for qRT-PCR in this study are: *Ehmt2*: 5’-ACCCCTCAGAGTGAGGAGAC-3’ (forward), 5’-GACGGTGACAGTGACAGAGG-3’ (reverse); *Runx2*: 5’-ATCCCCATCCATCCACTCCA-3’ (forward), 5’-GCCAGAGGCAGAAGTCAGAG-3’ (reverse); *Gapdh*: 5’-TGAAGGTCGGTGTGAACGG-3’ (forward), 5’-CAATCTCCACTTTGCCACTGC-3’ (reverse).

### Investigating *Ehmt2* and *Ehmt1* expression level from single cell data set

The mouse limb single-cell RNA-seq data set from the ENCODE Consortium mouse embryo project ^18^ was used to investigate *Ehmt2* and *Ehmt1* expression level across 25 cell types defined previously in E15 mouse limb tissue. The processed 10 X Genomics data was downloaded at https://cells.ucsc.edu/mouse-limb/10x/200120_10x.h5ad, SeuratDisk v0.0.0.9015 was used to convert the h5ad format into a Seurat object. R package dittoSeq v1.1.0 was used to visualize *Ehmt2* and *Ehmt1* mRNA expression levels across all cell types with box plots.

## Results

### Ehmt2 ablation in neural crest cells disrupts craniofacial morphogenesis

To investigate the role of *Ehmt2* in neural crest cell (NCC)-derived skeletal development, we generated *Wnt1-Cre; Ehmt2*^*f/f*^ mice. We found that the *Wnt1-Cre; Ehmt2*^*f/f*^ mice exhibited postnatal growth failure and craniofacial abnormalities. Micro-computed tomography (µCT) analysis of E17.5 embryos revealed microcephaly and a partial delay in the formation of the nasal, frontal, and parietal bones (**Fig. 1A**). The skeleton was also stained with Alcian Blue and Alizarin Red S to visualize the cranial morphology (**Fig. 1B**). *Wnt1-Cre; Ehmt2*^*f/f*^ embryos displayed impaired chondrogenic differentiation in the nasal region and significant malformations of the lower jaw, including a shortened and thickened mandible compared to controls (WT/*Ehmt2*^*f/f*^). The paranasal cartilage (PNC), premaxillary (PX), and maxillary bones were also severely affected. Moreover, ossification of the basisphenoid (BS) and basioccipital (BO) bones was markedly reduced, and the spheno-occipital synchondrosis (SOS) was abnormally widened, indicating disrupted cranial base development. Collectively, these findings demonstrate that *Ehmt2* is essential for proper craniofacial morphogenesis and jaw formation in NCC-derived tissues.

**Figure 1.**
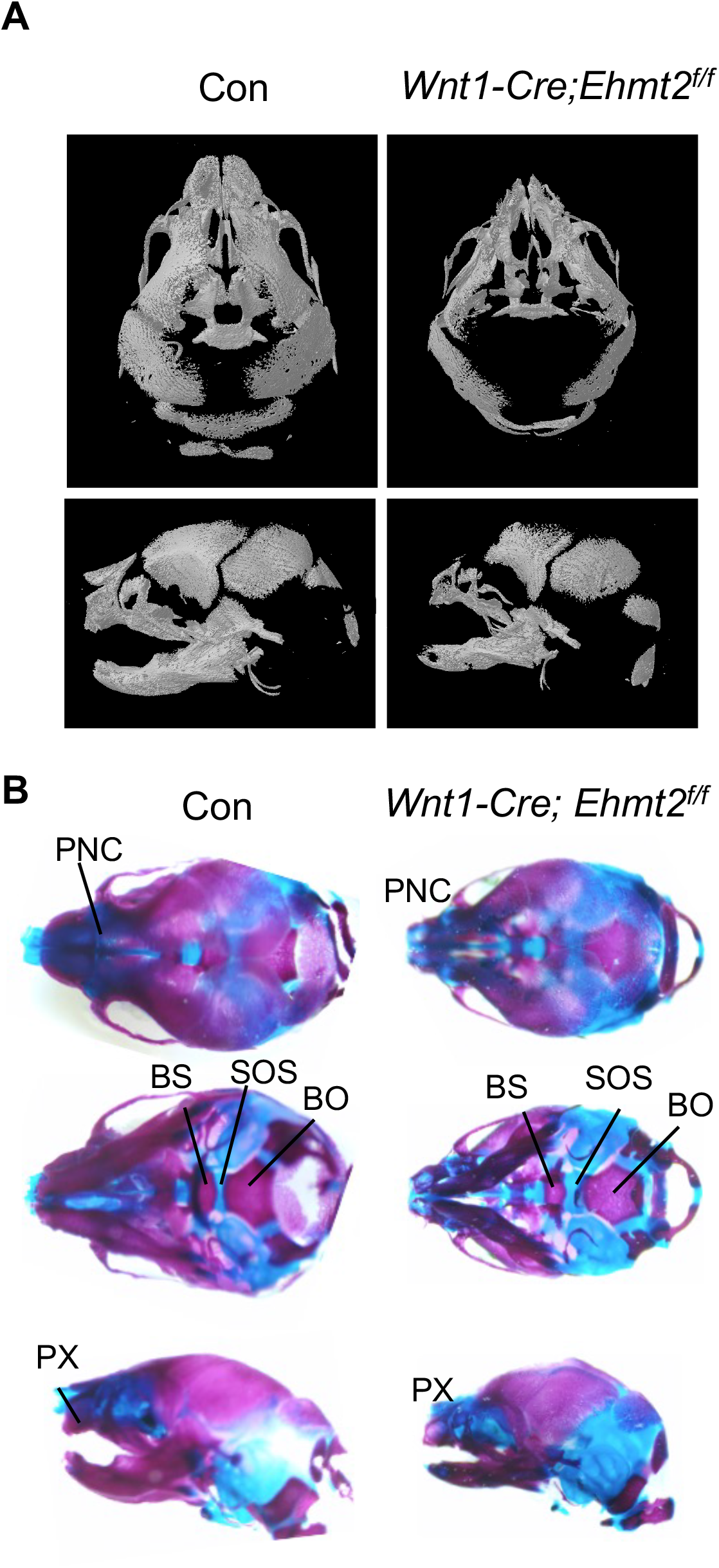
*Ehmt2* ablation in neural crest cells causes craniofacial hypoplasia and cranial base defects. (**A**) Micro-computed tomography (µCT) analysis of E17.5 embryos revealed microcephaly and delayed ossification of the nasal, frontal, and parietal bones in *Wnt1-Cre; Ehmt2*^*f/f*^ embryos compared to controls (*Ehmt2*^*f/f*^). (**B**) Alcian Blue and Alizarin Red S staining of whole-mount skeletal preparations highlighted defects in cranial morphology. *Wnt1-Cre; Ehmt2*^*f/f*^ embryos exhibited impaired chondrogenic differentiation in the nasal region and severe mandibular malformations, including a shortened and thickened lower jaw. The paranasal cartilage (PNC), premaxillary (PX), and maxillary bones were also significantly affected. Furthermore, ossification of the basisphenoid (BS) and basioccipital (BO) bones was markedly reduced, and the spheno-occipital synchondrosis (SOS) appeared abnormally widened, indicating disrupted cranial base development.

### Ehmt2 Loss Disrupts Proliferation and Activates Inflammatory Pathways in Neural Crest Cells

To investigate the molecular alterations associated with the craniofacial defects observed in *Wnt1-Cre; Ehmt2*^*f/f*^ mice (KO), we performed RNA sequencing on NCCs isolated from E10.5 embryos of both KO and control (*Ehmt2*^*f/f*^, WT) genotypes (n=2 per group), a developmental stage when neural crest cells are actively migrating and contributing to craniofacial structures. Unsupervised hierarchical clustering (Euclidean distance; **Fig. 2A**) and principal component analysis (PCA; **Fig. 2B**) demonstrated transcriptional divergence between the KO and WT NCCs. PC1 account for 83% of the variance (**Fig. 2B**), indicating widespread *Ehmt2*-dependent gene expression changes. Differential gene expression analysis identified a set of significantly upregulated and downregulated genes in KO NCCs compared to WT NCCs (**Fig. 2C, Supplemental Table 1**). Although *Ehmt2* is a known transcriptional repressor via H3K9me2 deposition, its loss in NCCs led to a global downregulation of gene expression, including genes encoding histone modifiers (*Setdb1, Suv39h1, Suv39h2, Hdac2, Ehmt1*), chromatin components (*H3c11, H3c13*), and RNA processing factors (*Celf1, Srsf1, Srsf6, Srsf9, Dicer1*). These findings suggest that *Ehmt2* plays a broader, non-canonical role in maintaining transcriptional and chromatin regulatory networks essential for NCC identity and proliferation.

**Figure 2.**
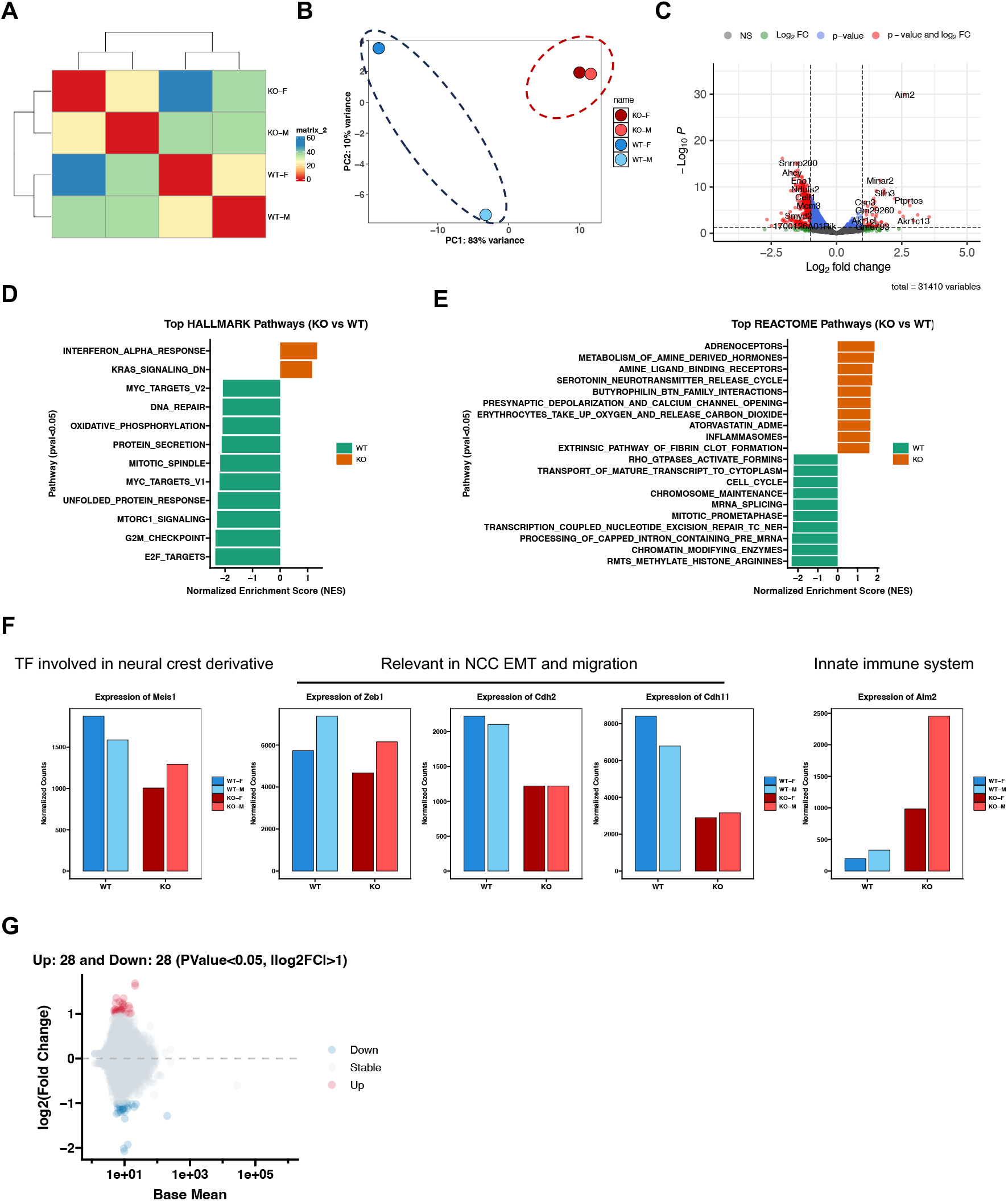
*Ehmt2* ablation alters transcriptional programs in neural crest cells. (**A**) Unsupervised hierarchical clustering of differentially expressed genes (DEGs) in *Wnt1-Cre; Ehmt2*^*f/f*^ (KO) vs. control (*Ehmt2*^f/f^, WT) NCCs (E10.5; n=2 per group). Euclidean distance demonstrates distinct transcriptional profiles. (**B**) Principal component analysis (PCA) confirms separation of KO and WT NCCs. (**C**) Volcano plot of DEGs (|log2FC| > 1 and p-value < 0.05). (**D**-**E**) Gene set enrichment analysis (GSEA) of Hallmark and REACTOME pathways. Top enriched terms in KO NCCs include inflammatory and neuromodulatory pathways, while WT NCCs exhibit proliferative and cell cycle signatures. NES: normalized enrichment score; FDR: false discovery rate. (**F**) Heatmap of select deregulated transcription factors (TFs) and NCC functional genes. *Meis1* and EMT/migration genes (*Zeb1, Cdh2, Cdh11*) are downregulated, while the innate immune gene *Aim2* is upregulated in KO NCCs. (**G**) MA plot showing differentially expressed TE loci in *Wnt1-Cre; Ehmt2*^*f/f*^ (KO) versus *Ehmt2*^*f/f*^ (WT) neural crest cells. TE expression was quantified using SQuIRE. Only intergenic and intronic TEs were included in the analysis. Significance thresholds were set at *p* < 0.05 and |log_2_ fold change| > 1.

Gene set enrichment analysis (GSEA) using Hallmark and REACTOME databases revealed that *Ehmt2*-deficient NCCs exhibited strong enrichment of inflammatory and neuromodulatory pathways, including interferon-alpha response, inflammasome activation, adrenoceptor signaling, and serotonin release (**Fig. 2D-E**). In contrast, WT NCCs were enriched for proliferative and cell cycle-related pathways, such as E2F targets, G2/M checkpoint, mTORC1 signaling, and chromosome maintenance. Further analysis of transcription factors and NCC lineage genes revealed a significant downregulation of *Meis1*, a key regulator of cranial NCC development (**Fig. 2F**). Genes involved in epithelial-to-mesenchymal transition (EMT) and migration, such as *Zeb1, Cdh2*, and *Cdh11*, were also suppressed. Conversely, upregulation of *Aim2*, a cytosolic double-stranded DNA sensor and mediator of innate immunity ^19,20^, points to heightened inflammatory signaling in the absence of *Ehmt2*.

Notably, *Ehmt2*, together with *Setdb1, Suv39h1/2*, and *Ehmt1*, are known to contribute to heterochromatin formation and the epigenetic repression of repetitive genomic regions–including transposable elements (TEs) and endogenous retrovirus (ERVs). We next examined whether its loss resulted in TE derepression/reactivation. However, global TE expression remained largely unchanged in *Ehmt2*-deficient NCC (**Fig. 2G)**. This paradox may be due to concurrent downregulation of cell cycle genes and transcriptional machinery, limiting TE activation, or to the compensatory chromatin silencing mechanisms in NCCs that maintain the heterochromatin occupied by TEs. Collectively, the chromatin-related changes indicate a loss of epigenetic stability, potentially contributing to the observed cycle arrest and inflammatory activation upon *Ehmt2* depletion.

### Ehmt2 is Dispensable for Skeletal Development in Osteochondral Progenitors

To determine whether the requirement for *Ehmt2* in skeletal development is lineage-specific, we next examined its role in *Prx1*-expressing osteochondral progenitor cells. Given that *Prx1-Cre* is expressed in mesenchymal progenitors of the limb and a subset of craniofacial regions ^14^, we generated *Prx1-Cre; Ehmt2*^*f/f*^ mice to investigate the role of *Ehmt2* in osteochondral progenitor cells. These mice were viable, fertile, and born at the expected Mendelian ratio, with no gross abnormalities in limb size or morphology. µCT of 1-month-old *Prx1-Cre; Ehmt2*^*f/f*^ mice showed no detectable differences in cranial bone mineralization compared to littermate controls (**Fig. 3A**). Similarly, analysis of trabecular and cortical bone parameters in the femurs of both sexes at 2 months of age showed no significant alterations (**Fig. 3B-D**).

**Figure 3.**
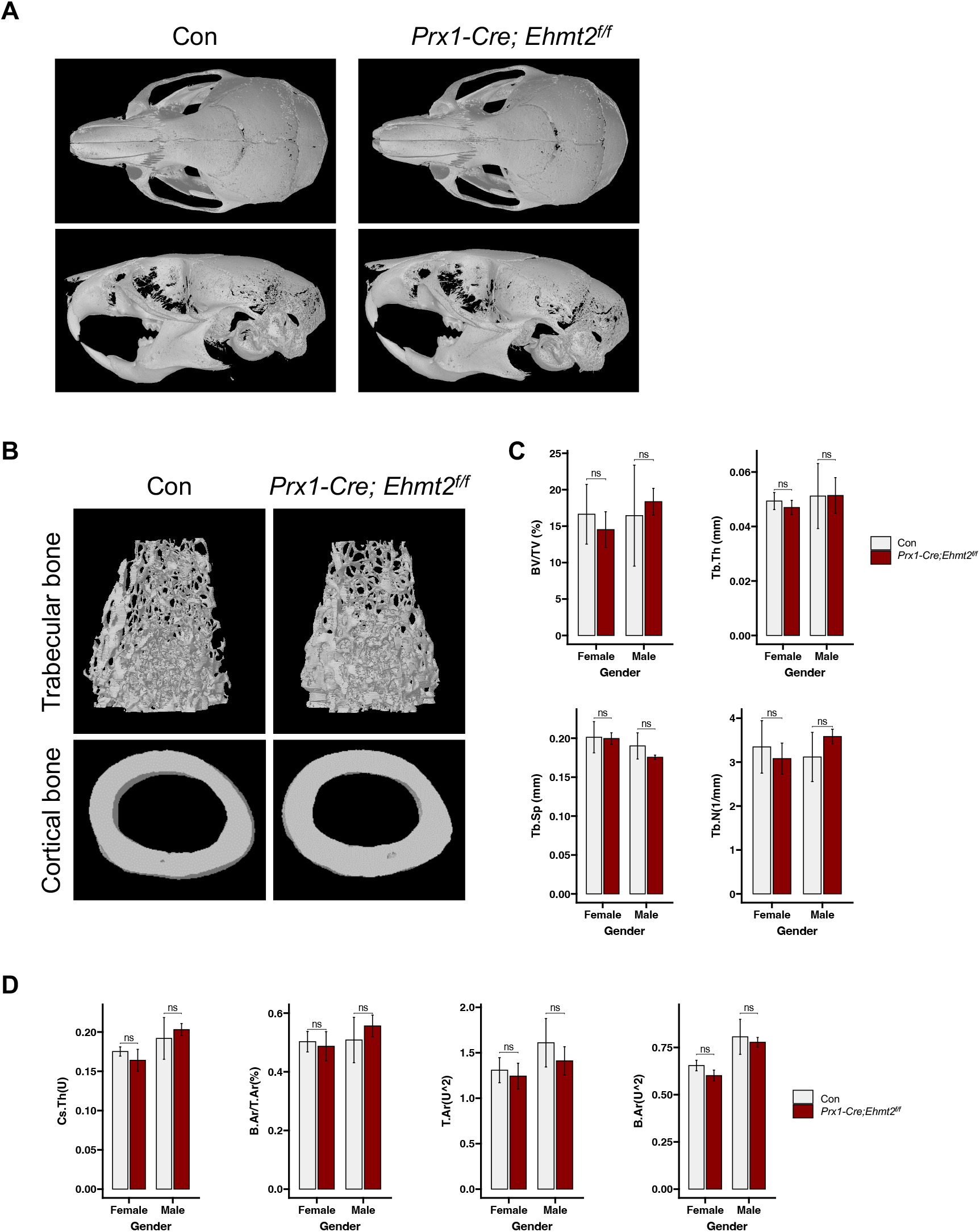
µCT analysis of cranial and femoral bones in *Prx1-Cre; Ehmt2*^*f/f*^ and *Ehmt2*^*f/f*^ control mice. (**A**) µCT imaging of the cranial skeleton shows no differences in bone morphology or mineralization between *Ehmt2*^*f/f*^ (Con) and *Prx1-Cre; Ehmt2*^*f/f*^ (*Ehmt2* cKO) mice at 1 month of age. (**B**) Representative µCT images of femoral trabecular bone (upper panels) and cortical bone (lower panels) from 2-month-old male Con and *Ehmt2* cKO mice. (**C**) Quantification of trabecular bone parameters of both sexes, including bone volume fraction (BV/TV), trabecular number (Tb.N), trabecular separation (Tb.Sp), and trabecular thickness (Tb.Th). (**D**) Cortical bone parameters, cortical thickness (Cs.Th), bone area/total area ratio (B.Ar/T.Ar), total cross-sectional area (T.Ar), and bone area (B.Ar) were compared between genotypes (ns = not significant; ^*^ *p* < 0.05; ^**^ *p* < 0.01; ^***^ *p* < 0.001).

To confirm *Ehmt2* gene deletion in *Prx1-Cre; Ehmt2*^*f/f*^ mice, we performed real-time quantitative PCR (qPCR) using RNA isolated from the femoral bone shafts at postnatal day 21. As expected, *Ehmt2* expression was significantly reduced in *Prx1-Cre; Ehmt2*^*f/f*^ mice compared with control (**Fig. 4A**). The expression of *Runx2*, a key master transcription factor for osteoblast differentiation, was not significantly altered (**Fig. 4A**). To further evaluate whether *Ehmt2* loss affects osteogenic potential, we conducted the *ex vivo* bone marrow stromal cells (BMSCs) differentiation assay. Alizarin Red staining of osteogenic cultures revealed reduced mineral deposition at day 8 in *Prx1-Cre; Ehmt2*^*f/f*^ MSC; however, this difference was largely resolved by day 14 (**Fig. 4B**), suggesting a delayed, but not blocked, osteogenic differentiation. To assess whether *Ehmt2* affects osteoblast function *in vivo*, we performed dynamic histomorphometry analysis. Calcein/alizarin double labeling of cortical midshaft (**Fig. 4C**) and distal femoral trabecular bone (**Fig. 4D**) revealed no significant differences in mineral apposition rate or bone formation parameters.

**Figure 4.**
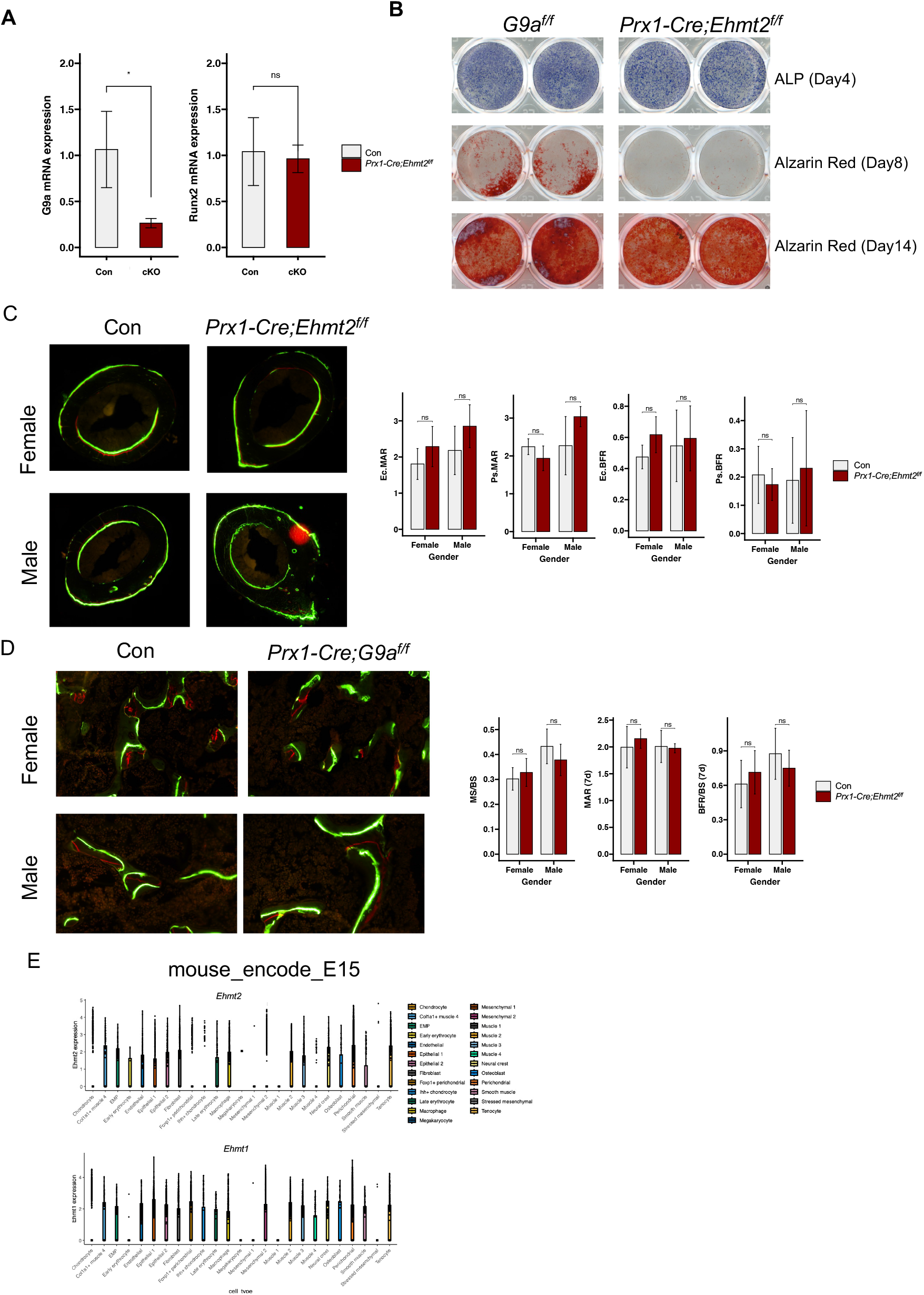
BMSC *ex vivo* osteogenic differentiation. (**A**) RT-qPCR analysis of relative *Ehmt2* and *Runx2* expression in 1-month-old *Ehmt2*^*f/f*^ (Con) and *Prx1-Cre; Ehmt2*^*f/f*^ (*Ehmt2* cKO) hindlimb bone osteoblasts. Statistical significance was assessed using Student’s t-test on Ct values (n = 3; ns = not significant; ^*^ *p* < 0.05; ^**^ *p* < 0.01; ^***^ *p* < 0.001). (**B**) *Ex vivo* osteogenic differentiation of BMSCs from 1-month-old male *Ehmt2*^*f/f*^ (Con) and *Prx1-Cre; Ehmt2*^*f/f*^ (*Ehmt2* cKO), cultured in 24-well plates. (**C**) Calcein/alizarin double labeling of the cortical midshaft of femurs from 2-month-old *Ehmt2*^*f/f*^ (Con) and *Prx1-Cre; Ehmt2*^*f/f*^ (*Ehmt2* cKO) mice was used to assess dynamic bone formation. No significant differences were observed in mineral apposition rate (MAR) or other formation parameters. (**D**) Calcein/alizarin labeling of trabecular bone in the distal femur also showed no significant differences in Mineral Apposition Rate (MAR) or bone formation rate between genotypes. (**E**) Publicly available ENCODE RNA-seq data from embryonic day 15 (E15) mouse tissues showed high expression of *Ehmt2* and *Ehmt1*in neural crest-derived tissues, but relatively low expression in limb mesenchymal cells. This differential expression may help explain the distinct skeletal phenotypes observed with tissue-specific deletion of *Ehmt2*.

To explore whether cell-type-specific expression of *Ehmt2* may underlie differential phenotypes, we analyzed publicly available ENCODE RNA-seq data from E15 mouse tissues. *Ehmt2* and *Ehmt1* were both highly expressed in neural crest derivatives, but relatively low in limb mesenchymal cells (**Fig. 4E**). This differential expression pattern may help explain why *Ehmt2* ablation in NCCs leads to profound craniofacial abnormalities, whereas its loss in *Prx1*-expressing mesenchymal lineages has minimal impact on skeletal development. Together, these results indicate that Ehmt2 may not be critically required for osteochondral progenitor function or bone formation under homeostatic conditions.

## Discussion

In this study, we investigated the role of the histone methyltransferase *Ehmt2* in skeletal development by conditionally deleting it in neural crest- and mesoderm-derived progenitors using *Wnt1-Cre* and *Prx1-Cre* mice. Our findings demonstrate that deletion of *Ehmt2* in cranial neural crest cells using the *Wnt1-Cre* transgene resulted in a distinct craniofacial phenotype in perinatal mice. µCT analysis and Alcian Blue and Alizarin Red S revealed delayed ossification in the nasal, frontal, and parietal bones, suggesting that *Ehmt2* is required for proper cranial bone development within the neural crest lineage. In contrast, deletion of *Ehmt2* in osteochondroprogenitor cells using the *Prx1-Cre* transgene did not lead to any overt abnormalities in limb morphology or size. µCT analysis showed no significant differences in trabecular or cortical bone parameters between *Prx1-Cre; Ehmt2*^*f/f*^ mice and control *Ehmt2*^*f/f*^ littermates.

Histomorphometry analyses further supported these findings, revealing no major differences in bone formation or cartilage morphology between the two groups. Despite the limited craniofacial activity of *Prx1-Cre* ^14^, we did not observe craniofacial phenotypes in *Prx1-Cre; Ehmt2*^*f/f*^ mice, suggesting that *Ehmt2* plays a minimal or redundant role in these regions compared to its function in neural crest-derived craniofacial progenitors.

From our RNA-seq analysis of NCCs, we found that *Ehmt2* knockout not only reduced the expression of *Ehmt1*—a known *Ehmt2* partner—but also led to a significant decrease in other H3K9 methyltransferases, including *Setdb1* and *Suv39h1/2*. This observation is consistent with previous findings in somatic cell nuclear transfer (SCNT) embryos, where pharmacological inhibition of *Ehmt2* led to a reduction in H3K9me1/2 levels and, unexpectedly, a decrease in H3K9me3 levels as well ^21^. Given that *Ehmt2* forms multimeric complexes with other H3K9 methyltransferases ^22^, these results suggest that *Ehmt2* loss may destabilize other components of the H3K9 methylation network, leading to broader epigenetic dysregulation. While H3K9 methylation is typically linked to transcriptional repression, we observed a substantial number of downregulated genes in *Ehmt2* knockout NCCs, highlighting the non-canonical and context-dependent roles of *Ehmt2* in gene activation. These results are consistent with prior studies showing that EHMT2–EHMT1 complexes regulate euchromatic H3K9me1/2 and support proper chromatin organization ^3^. Disruption of this complex can lead to global transcriptional changes, including HP1 relocalization and chromatin disorganization.

Despite *Ehmt2*’s well-established role in TE silencing, our transcriptomic analysis revealed no significant increase in TE expression upon *Ehmt2* loss in E10.5 NCCs. This may reflect compensatory activity from partially retained *Setdb1*/*Suv39h1/2* expression, or involvement of alternative silencing pathways such as DNA methylation or PRC2-mediated H3K27 methylation ^23^. It is also likely that the concurrent downregulation of cell cycle regulators and components of the transcriptional machinery limits TE activation. Additionally, TE derepression may require stress-related signals or chromatin priming events that are absent during early development before NCCs differentiate.

Although *ex vivo* osteogenic differentiation of mesenchymal stem cells (MSCs) from *Prx1-Cre; Ehmt2*^*f/f*^ mice showed a transient and moderate reduction in mineral deposition at day 8, this difference was largely resolved by day 14, indicating only a mild and temporary defect in osteogenesis under in vitro conditions. In contrast, *in vivo* analysis by µCT and dynamic histomorphometry revealed no significant changes in bone mass or formation rate. Therefore, while *Ehmt2* loss may initially delay osteogenic commitment in a simplified in vitro environment, these effects appear to be compensated or mitigated in vivo, suggesting that *Ehmt2* is not critically required for osteoblast differentiation and bone formation under normal physiological conditions.

In conclusion, these results underscore a differential requirement for *Ehmt2* in distinct skeletal progenitor lineages and highlight its critical role in neural crest-derived craniofacial development and provide molecular insights for the human neurodevelopmental and craniofacial diseases, such as KS1 and KS-like syndromes.

## Acknowledgments

We thank the current and former members of Dr. T. Yang’s laboratories for their contributions, as well as B. Williams, for discussions, and D. Brass for editing. This work was supported by the dedicated staff of VAI Vivarium (RRID:SCR_023211), Genomics (RRID:SCR_022913), and Pathology & Biorepository Cores at VAI (RRID:SCR_022912).

## Data availability

Data will be made available on request.

## Author contributions

Y.L. and T.Y. conceived this study and designed experiments. Y.L., Y.Z., P.K., M.L., J.L., H.L., and D.L. executed experiments and performed data analysis. Y.L., H.L., and P.K. generated figures. Y.L. wrote the manuscript, and T.Y., M.L, and P.E.S. edited the manuscript.

